# Multiscale simulations elucidate the mechanism of polyglutamine aggregation and the role of flanking domains in fibril polymorphism

**DOI:** 10.1101/2025.05.19.654960

**Authors:** Avijeet Kulshrestha, Tien Minh Phan, Azamat Rizuan, Priyesh Mohanty, Jeetain Mittal

## Abstract

Protein aggregation, which is implicated in aging and neurodegenerative diseases, typically involves a transition from soluble monomers and oligomers to insoluble fibrils. Polyglutamine (polyQ) tracts in proteins can form amyloid fibrils, which are linked to polyQ diseases, including Huntington’s disease (HD), where the length of the polyQ tract inversely correlates with the age of onset. Despite significant research on the mechanisms of Httex1 aggregation, atomistic information regarding the intermediate stages of its fibrillation and the morphological characteristics of the end-state amyloid fibrils remains limited. Recently, molecular dynamics (MD) simulations based on a hybrid multistate structure-based model, Multi-eGO, have shown promise in capturing the kinetics and mechanism of amyloid fibrillation with high computational efficiency while achieving qualitative agreement with experiments. Here, we utilize the multi-eGO simulation methodology to study the mechanism and kinetics of polyQ fibrillation and the effect of the N17 flanking domain of Huntingtin protein. Aggregation simulations of polyQ produced highly heterogeneous amyloid fibrils with variable-width branched morphologies by incorporating combinations of β-turn, β-arc, and β-strand structures, while the presence of the N17 flanking domain reduces amyloid fibril heterogeneity by favoring β-strand conformations. Our simulations reveal that the presence of N17 domain enhanced aggregation kinetics by promoting the formation of large, structurally stable oligomers. Furthermore, the early-stage aggregation process involves two distinct mechanisms: backbone interactions driving β-sheet formation and side-chain interdigitation. Overall, our study provides detailed insights into fibrillation kinetics, mechanisms, and end-state polymorphism associated with Httex1 amyloid aggregation.

**SIGNIFICANCE STATEMENT:** Polyglutamine (polyQ) aggregation is central to Huntington’s disease and related neurodegenerative disorders. Despite extensive experimental efforts, a complete molecular understanding of this process—from early aggregation events to the origins of fibril polymorphism—has remained elusive, with varied interpretations of complex fibril architectures. Through multiscale simulations, we reveal how polyQ fibrils adopt diverse tertiary and quaternary structures and demonstrate how the N-terminal flanking domain (N17) modulates fibril architecture and accelerates aggregation. Our hybrid multi-eGO simulations capture early-stage fibrillation kinetics and identify distinct structural polymorphs that align with experimental observations. This work provides a molecular framework for understanding amyloid polymorphism and illuminates the role of flanking domains in shaping aggregation pathways— offering valuable insights for therapeutic strategies targeting early toxic intermediates.

## INTRODUCTION

Polyglutamine (polyQ) tracts in proteins, encoded by CAG trinucleotide repeats, can form amyloid fibrils implicated in nine neurodegenerative diseases, each characterized by a critical threshold of polyQ (CAG) expansion beyond which tract length correlates with age-at-onset and clinical severity (1). The exon 1 of Huntingtin (Htt) protein (2) is intrinsically disordered, containing a central polyQ tract whose length varies between 10 and 24 glutamine (Q) residues in the wild-type protein. The expansion of the polyQ tract length in Htt (>Q35) is associated with Huntington’s disease (HD) (2, 3). The flanking domains in Httex1 include a 17 residue N-terminal coiled-coil domain (N17) and 51 residue proline-rich domain (PRD) C-terminal domain. The N17 domain has previously been shown to greatly enhance the aggregation kinetics of Httex1 in vitro (4) through the initial formation of soluble oligomers which increases the spatial proximity of polyQ tracts (4, 5). Importantly, the expression of Httex1 alone is sufficient to recapitulate the HD phenotype *in vivo* (6). The structural ensembles of Httex1 wild-type and pathogenic polyQ length have been studied extensively with solution NMR (5, 7) and SAXS (8). These investigations reveal an emergent propensity for more favorable α-helical structure formation in the polyQ tract with increasing length from wild type to near and beyond the pathological limit.

Unlike other well-studied amyloid forming proteins, atomic-resolution structures of polyQ and Httex1 amyloid fibrils has remained inaccessible by modern cryo-EM methods due to high structural heterogeneity associated with polyQ self-assembly and the conformational disorder associated with the flanking regions (5, 9). Low-resolution structural investigations using negative stain transmission electron microscopy (TEM) and atomic force microscopy (AFM) both show that polyQ peptide and HttEx1 produce extensively branched filament-type assemblies, where the kinetics of the self-assembly depends not only on the polyQ length but also on the presence of N17, as observed with HPLC-based sedimentation assays (4, 10–12). A recent cryo-EM study revealed structural polymorphism within protofilament is composed of the stacks of β-hairpins and β-strand which show variable stacking angles with occasional out-of-register states. Interestingly, removal of the N-terminus results in higher intrafilament structural heterogeneity, which suggests that flanking domains appear to play an important role in stabilizing the polyQ core by preventing large out-of-register shifts (13, 14). Another recent integrative approach combining solid-state (ss) NMR and molecular dynamics was used to determine one possible core structure of polyQ and Huntington exon 1 (HttEx1), where the polyQ fibril core of HttEx1-Q44 was modeled with a single β-turn (15). The structure features of polyQ fibrils vary considerably across reported studies (7, 13, 16) and overall, the HttEx1 fibril exhibits both supramolecular and protofilament polymorphism (17).

Despite extensive research efforts to understand the process of HttEx1 amyloid formation, a complete molecular view of the events involving the soluble monomers into amyloid fibrils is currently unattainable using experimental approaches alone. With modern GPU-based hardware and algorithms, it is now feasible to perform multi-microsecond timescale simulations of protein folding and other conformational transitions in explicit solvent using classical all-atom molecular dynamics (AAMD) simulations. Brute-force AAMD in combination with enhanced sampling methods have been extensively employed to study the polyQ length-dependent structure and conformational transitions of Huntingtin (18–20). However, self-assembly processes which occurs during protein aggregation such as nucleation and elongation which occur on the timescale of several milliseconds to hours and days remain inaccessible to AAMD at present. As an alternative to expensive AAMD simulations, Wolynes and co-workers simulated the free energy landscapes of polyglutamine aggregation computed using an efficient, three-site per amino acid coarse-grained (CG) protein model (21) In line with the experimental observations of Wetzel and co-workers, CG simulations demonstrated that shorter polyQ_20_ monomer prefers an extended conformation and a trimeric nucleus (n*∼3), while the longer Q_30_ preferably adopts a β-hairpin conformation and aggregates through a monomeric nucleus (n*∼1). Alternatively, Schmit and co-workers investigated polyQ thermodynamics using triblock co-polymer models of HttEx1 to explain oligomer structure and flanking sequence effects, and AAMD-derived lattice models with binary amino sequence states to show how conformational entropy governs β-sheet nucleation and elongation (22, 23).

Camilloni and co-workers recently developed a hybrid multistate structure-based model– Multi-eGO, which enables the study of protein aggregation in feasible simulation time at the level of heavy atom resolution for the polypeptide in implicit solvent (24, 25). Multi-eGO enables a concentration-dependent assessment of the fibrillation kinetics and mechanisms which are shown to qualitatively agree with experiments for small peptides (26, 27). Here, we parameterized multi-eGO models to investigate the fibrillation kinetics and mechanism of polyQ amyloid aggregation. Multi-eGO aggregation simulations of Q16 revealed a high level of heterogeneity at the protofilament level, consisting of β-turn, β-arc, and β-strand structures. In contrast, the addition of an N-terminal coiled-coil domain (N17) and C-terminal polyproline motif (P_5_) from Httex1 to Q16 gave rise to protofibrils with a higher propensity of β-strand formation. We further analyzed the secondary structure content in protofibrils taken from the multi-eGO end-state fibril using explicit solvent AA simulations and observed a partial helicity in the N-terminus with a stable polyQ core arrangement. Interestingly, our multi-eGO simulations revealed that the presence of N17 domain enhanced aggregation kinetics by promoting the formation of large, structurally stable oligomers. Moreover, this study reveals distinct pathways in early-stage aggregation, one involving the establishment of ordered β-sheet architecture via key backbone interactions and another facilitated by the cooperative interlocking of side chains. In summary, our results provide rich insights into the fibrillation mechanism, end-state fibril heterogeneity, and the effect of N17 domain on the aggregation kinetics of Httex1.

## RESULTS AND DISCUSSION

The multi-eGO method integrates native contact information obtained from all-atom simulations for individual free-energy minima of the system. For a system undergoing fibrillation, these minima correspond to the native soluble and fibril states, respectively. This allows for simulating the aggregation process from monomers to fibrils, using contact information obtained from the two states to capture intermediate oligomeric states.

Based on *in vivo* Distributed Amphifluoric FRET (DAmFRET) and MD simulations, Halfman and co-workers (28) determined that polyQ amyloid formation begins with the formation of a minimal steric zipper of six interdigitated sidechains (“Q-zipper”) within a single polyQ chain exceeding the pathological threshold. Considering the pathological threshold of Q36 for Huntington’s disease, such an arrangement requires six Qs per zipper strand and a minimum of four residues for modeling a turn between two strands (28), resulting in a total of 16 Qs per steric zipper unit. Further, our previous atomistic simulations validated against solution NMR showed that Httex1 constructs with a polyQ length of 16 residues (N17-Q16-P5) accurately capture essential conformational features of the polyQ tract, including helical propensities and domain interactions relevant to early aggregation events (18). An accurate all-atom single-chain ensemble is crucial for multi-eGO contact parameterization of the monomeric state. Therefore, we selected a 16-residue polyQ tract for our study which satisfies both minimal steric zipper requirements and can be validated against an accurate all-atom conformational ensemble, for multi-eGO model parameterization.

### Parameterization of monomer contacts for multi-eGO

To develop multi-eGO models of Q16 and N17-Q16-P_5_ (H16), we first parameterized the intramolecular contacts expected to form in the monomeric state. The monomer multi-eGO models were parameterized to reproduce intramolecular (non-bonded) interactions formed in all-atom (AA) monomer ensembles generated from explicit solvent MD simulations. CD and NMR experiments indicate that polyglutamine peptides largely populate a random coil ensemble in solution (29, 30), while H16 populates a partial α-helical structure (31). To obtain realistic structural ensembles of Q16/H16 in soluble state, we performed one microsecond-long AA simulations of both monomers using the AMBER03ws force field to extract the native contact pairs formed in these ensembles (see Methods). This force field was demonstrated to generate structural ensembles of Q16/H16 found to be in excellent agreement with NMR experiments (18).

The multi-GO model requires system-specific parameterization of the reference interaction strength (ε) of native contacts. For this study, ε was optimized to 0.4 kJ/mol for Q16 and 0.325 kJ/mol for H16, enabling the multi-eGO ensembles to reproduce key structural observables from AAMD simulations (**Figure S1**). Specifically, the multi-eGO radius of gyration (R_g_) distributions for both Q16 and H16 showed mean values and peak positions similar to those from AAMD **(Figure B, E)**, albeit with a slightly narrower distribution for the multi-eGO ensemble. Pairwise intramolecular contact probability maps (**Figure C**, **F)** also demonstrated strong agreement between AAMD and multi-eGO models. In terms of secondary structure, the multi-eGO simulations successfully captured the predominantly random coil nature of Q16 and the residual α-helical profile of H16 (**Figure D**, **G)**. Notably, both AA and multi-eGO simulations accurately reproduced the per-residue α-helical fractions of H16 obtained from NMR measurements (31, 32). Overall, the multi-eGO models capture the transient secondary structure and intramolecular interactions for both peptides, allowing us to proceed further to parameterize the fibril state.

### Parameterization of polyQ fibril state to drive amyloid aggregation from monomers using multi-eGO

While the atomistic details of the polyQ fibrils are not directly accessible to structure determination techniques, structural constraints inferred from X-ray diffraction and ssNMR (33) indicate that polyQ fibrils are characterized by a block-like, waterless amyloid core composed of multiple layers of tightly packed anti-parallel β-sheets. It is possible to arrange the antiparallel β-sheets in different secondary and quaternary configurations while following the experimental constraints and thereby account for the structural polymorphism associated with polyQ fibrils. Accordingly, we prepared the model Q16 fibril structures based on two possible tertiary structure units while satisfying the requirements of the minimal Q-zipper unit (28) - β-turn (BT) and β-arc (BA) (34). The β-turn model is characterized by a β-sheet structure in the lateral direction within a chain, and the interdigitation of side chains occurs with a different chain in the axial direction (**Figure 2A**). On the contrary, the interdigitation of side chains occurs within the molecule in the axial direction of the β-arc model, and the β-sheet is formed between strands of two different chains (**Figure 2E**).

**Figure 1:**
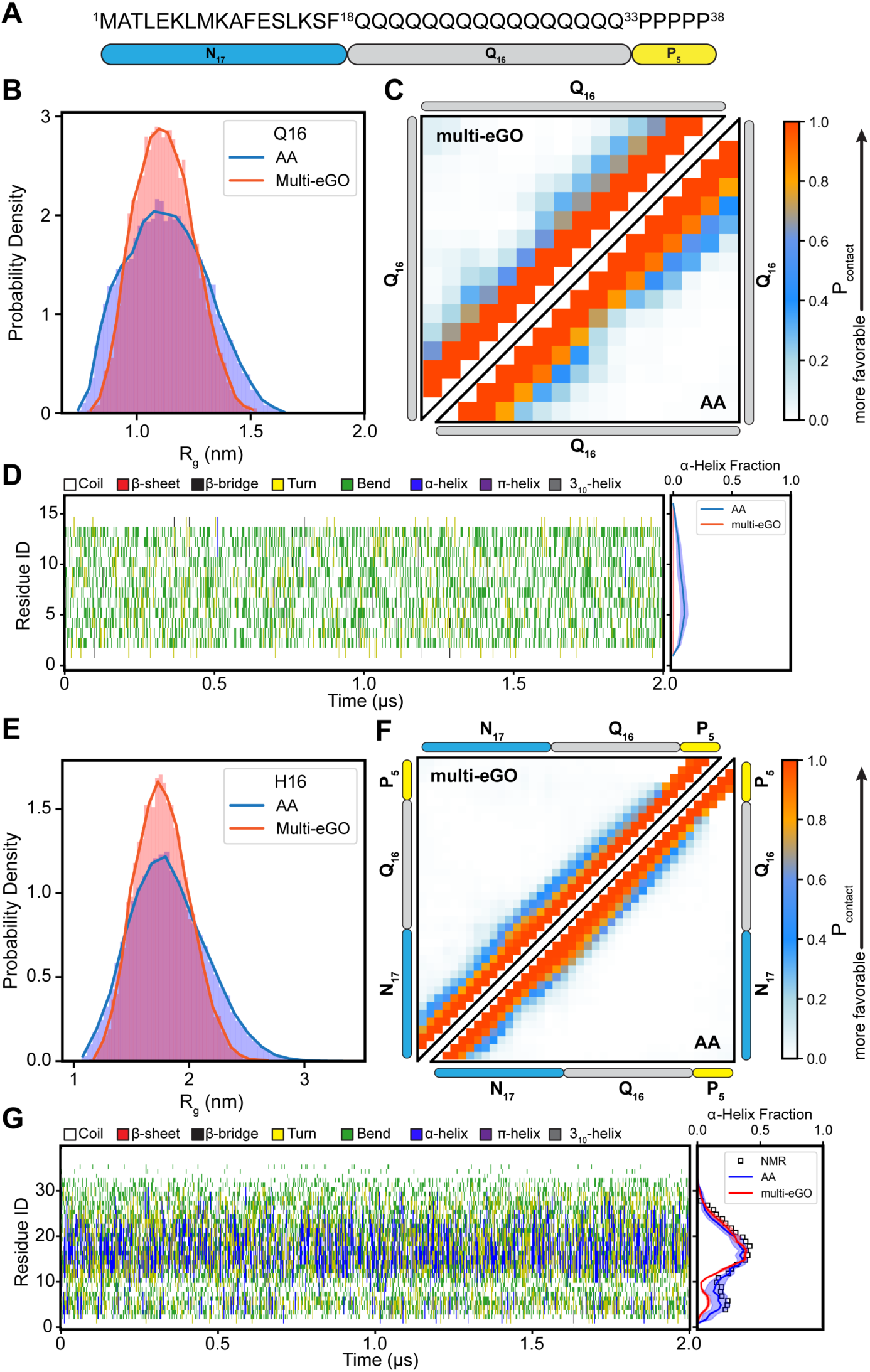
Multi-eGo modeling of Q16 and H16 monomer ensembles. (A) Schematic representation of the HttEx1 sequence, comprising the N-terminal N17 and the C-terminal P5 domains, flanking a central segment of 16 glutamine residues (Q16). (B) Radius of gyration distribution, (C) residue-based contact maps, and (D) time-dependent secondary structure variations for the Q16 peptide. (E) Radius of gyration distribution, (F) residue-based contact maps, and (G) time-dependent secondary structure variations for the H16 protein. Results from multi-eGO simulations are compared with corresponding all-atom simulation data. In the contact maps (panels C, F), the upper triangular region displays contacts from the multi-eGO simulations, while the lower triangular region shows contact data from all-atom simulations. The right panels in (D) and (G) shows the residual α-helix fraction of Q16 and H16, respectively. H16 helical fraction was compared to SSP scores for the helical propensity (square) computed from experimental 13C chemical shifts taken from Urbanek et al. (31, 32).

**Figure 2:**
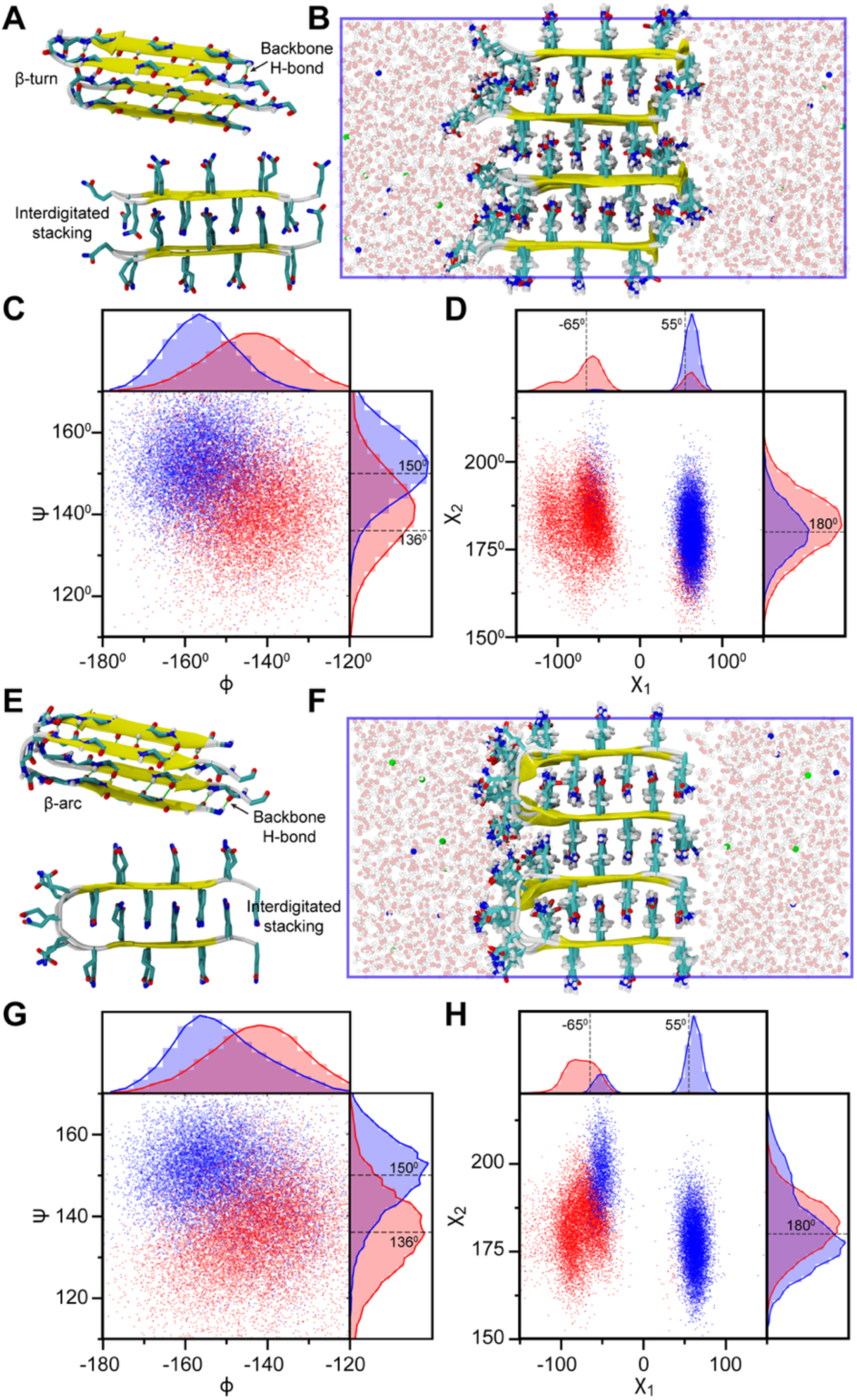
Structural characterization of two polyglutamine (PolyQ) fibril models in explicit-solvent AAMD simulations. (A) Top and side views of Q16 β-turn (BT) model, illustrating lateral backbone hydrogen bonds (green lines) and interdigitated sidechain stacking along the axial direction. (B) Semi-infinite simulation setup for the β-turn model. (C, D) Backbone and sidechain dihedral angle distributions for the β-turn model. (E) Top and side views of the β-arc (BA) model, highlighting analogous backbone hydrogen bonding and sidechain interactions. (F) Semi-infinite simulation setup for the β-arc model. (G, H) Backbone and sidechain dihedral angle distributions for the β-arc model. In (C, D, G, and H), red and blue represent dihedral angle values for alternating “a” and “b” conformer strands, and gray lines correspond to experimental ssNMR-derived values. Two separate lines indicate distinct dihedral values for each strand, whereas a single line denotes identical dihedral angle values observed in ssNMR for both strands. In (B, F), protein chains are depicted as ribbons, sidechains as licorice representation, highlighting interdigitated stacking, Na^+^ and Cl^-^ ions as blue and red spheres, respectively, and water molecules in ball-and-stick representation in the background.

To prepare stable quaternary fibril structures based on BT and BA monomer units, we arranged them in a semi-infinite cubic box, where we allow the periodic images to interact in X and Y directions (**Figures 2B, F**). Semi-infinite simulations of Alzheimer’s β-amyloid protofilament have previously been shown to reproduce the NMR-derived structural constraints (35, 36). We arranged the end-termini of two consecutive chains in four possible configurations for both BT and BA models (**Figure S2**): (i) both facing the same direction along the lateral axis (A1), (ii) oppositely in the lateral axis (A2), (iii) oppositely in the axial direction (A3), and (iv) oppositely in both lateral and axial directions (A4). To assess the stability of our fibril configurations, we first computed the average sheet-to-sheet and strand-to-strand distances from 1 μs trajectories for all directional arrangements of Q16 fibrils. The computed distances for all arrangements are in line with X-ray diffraction data (33), as shown in **Figure S3**. A higher distance variation is observed between two different chains, i.e., the variation in the sheet-to-sheet distance is higher for β-turn setups, and the strand-to-strand distance variation is higher for β-arc setups.

ssNMR studies of polyQ fibrils exhibit a characteristic 2D spectrum comprising of two sets of peaks which correspond to two dominant Q residue conformations—“a” and “b” conformers (33). The two Q conformers belong to two-distinct β-strands in an antiparallel β-sheet and differ in their ψ and χ_1_ dihedral angles. Therefore, we further validated our fibril arrangements against ssNMR-derived structural constraints comprising of ψ, χ_1_ and χ_2_ dihedral angles for both “a” and “b” conformers. We computed the same angles from the last 500 ns of MD trajectories for each alternative “a” and “b” strands separately for the A1 fibril arrangement (**Figure 2B, F**). AAMD simulations show two distinct distributions for “a” and “b” strands with peaks close to the ssNMR-derived values at 136 and 150° respectively (**Figures 2C, G**). Similarly, the χ_1_ angles of the AAMD distributions (**Figure 2D, H**) also exhibit major peaks near the experimental values at −65 and 55° for “a” and “b” strands respectively. Interestingly, in AAMD simulations, it was observed that each strand also sampled χ_1_ values associated with the other to a lesser extent, and such bimodal distributions were found to be more prominent for β-turn models than β-arc. Finally, χ_2_ angles distributions show a single peak near 180° that agrees with ssNMR for both models (**Figure 2D, H**). A similar analysis carried out for the remaining three arrangements of β-turn (**Figure S4**) and β-arc (**Figure S5**) models indicate that except for BT-A4, all other models also exhibit ψ, χ_1_ and χ_2_ dihedral angle distributions which are consistent with experiment.

In conclusion, our analysis demonstrates that polyQ fibril chains can adopt multiple packing arrangements consistent with ssNMR and X-ray diffraction measurements, highlighting the potential for molecular polymorphism at the secondary (turn vs. arc) and quaternary (directional arrangement) structural levels (33, 37–39). To investigate Q16 fibrillation, we developed two distinct multi-eGO models by training the fibril state with native contacts of β-turn and β-arc conformations obtained from the semi-infinite AAMD simulations in explicit water. In both models, fibril contacts were parameterized based on the A1 arrangements, in which all termini are oriented in the same directions (**Figure 2F**). The interaction strength (ε) for intermolecular contacts was set equal to that of intramolecular contacts, as this parameterization yielded structurally stable fibrils in the simulations.

### PolyQ amyloid fibrils exhibits the supramolecular polymorphism

Employing our multi-eGO models for Q16, which integrate contact information from monomeric states and the fibrillar structures based on two different tertiary units (BT/BA), we first performed aggregation simulations of 1000 chains for Q16 at a concentration of 10 mM at 300 K temperature. Three independent aggregation simulations were performed for each model until nearly all chains were present within the aggregate. Analysis of the total β-sheet fraction suggests that both β-turn and β-arc models show a similar behavior, and all the chains convert to β-sheet within 100 ns (**Figure 3A**). Furthermore, these multi-eGO simulations consistently generated variable-width, branched polyQ fibril morphologies (**Figure 3B**), which are strikingly similar to those observed in experimental studies using atomic-force, negative stain transmission electron and cryo-EM microscopy (4, 11, 13).

**Figure 3:**
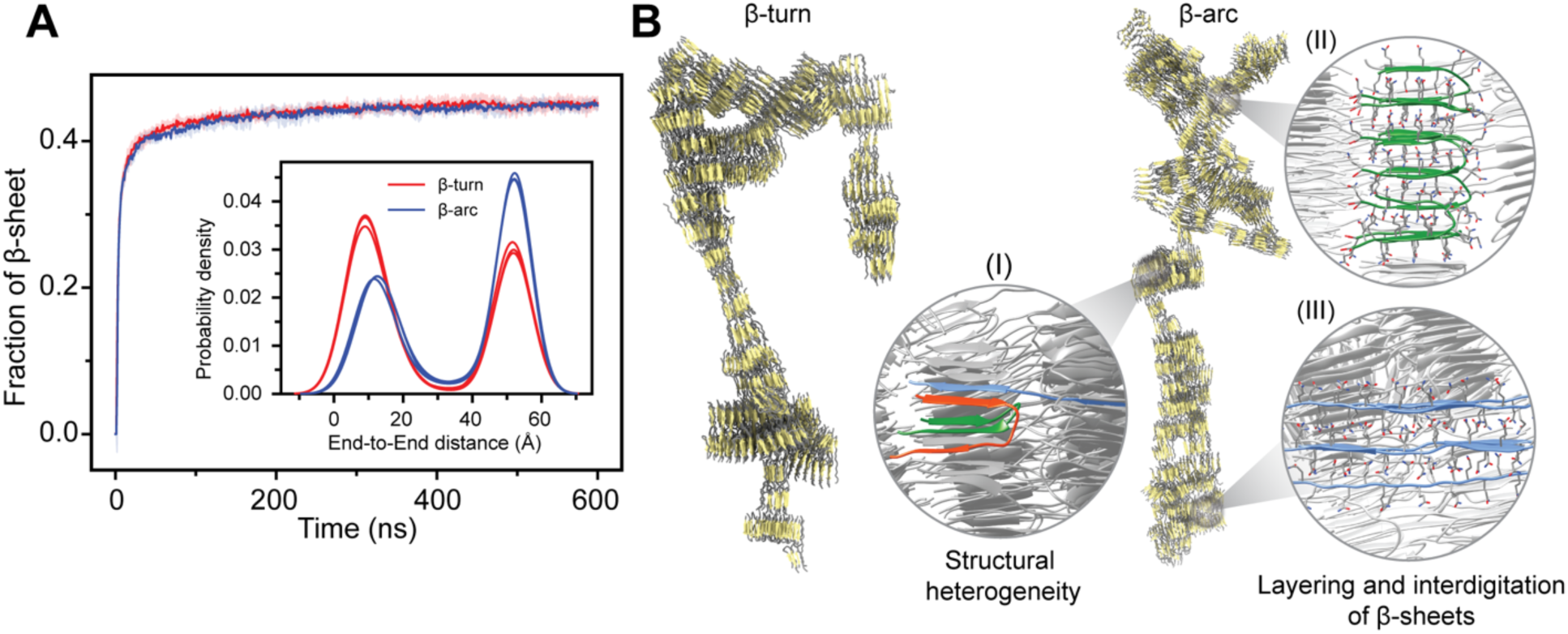
Structural heterogeneity of Q16 in multi-eGO aggregation simulations. (A) Time evolution of β-sheet fraction for Q16 β-turn and β-arc models simulated at 10 mM protein concentration and 300 K. The inset shows the corresponding end-to-end distance distribution, highlighting the coexistence of compact and extended conformations in the simulations. (B) Representative snapshots of Q16 β-turn and β-arc fibril. Enlarged snapshots of local fibrillar regions highlighting key structural details: (I) conformational polymorphism, showing the coexistence of β-turn (green), β-arc (red), and extended β-strand (blue) segments and (II) the characteristic layering and interdigitation of β-sheets. The overall branched morphology was formed by (III) extended β-strands that span between and connect different protofibrils.

Beyond corroborating the overall branched morphologies observed in our simulations, these cryoEM studies (4, 11, 13) further revealed a high degree of fibril polymorphism at the protofilament level which can be composed of the stacks of β-hairpins and linear β-strand that combine at variable stacking angles and occasional out-of-register positioning. For a quantitative assessment of structural heterogeneity at the protofilament level, we analyzed the end-to-end distance distribution of each chain from the final amyloid fibrils **(Figure 3A, inset)**, where a higher value indicates the extended β-strand configuration, and a small value indicates a compact structure composed of either β-turn or β-arc. Our analysis indicates that the polyQ chains in the amyloid fibril can exist in diverse configurations such as β-turn, β-arc and β-strand (extended). The β-arc multi-eGO model favors β-strand configurations over the compact structure, whereas β-turn model favors compact configurations. Such a preference for heterogeneous configurations in polyQ fibrils has been proposed to promote branched chain morphologies (40). Closer inspection of the fibril structure revealed that individual chains adopting β-strand configurations, can bridge multiple protofibrils, creating a branched morphology (**Figure 3B**). The axial interdigitation of side chains is a key characteristic of these inter-protofibril connections as well as the packing within the protofibrils themselves (**Figure 3B**). Overall, a similar degree of conformational heterogeneity at the level of tertiary structure and protofilament arrangement was observed regardless of whether the multi-eGO model was trained with contact information from the β-turn or β-arc fibril structure. While the ability of multi-eGO to sample a wide range of states likely stems from multiple aspects of its design, these findings indicate that the transferable bonded interactions within its framework are a contributing factor to exploring conformations beyond those directly encoded by the initial contact information. Therefore, multi-eGO models offer a pathway to identify conformational possibilities not explicitly defined in their initial parameterization.

### Parameterization of N17 domain intermolecular interactions for Multi-eGO

PolyQ proteins contain flanking sequences which play critical roles in modulating oligomerization and fibrillation. For example, Httex1 includes an N-terminal coiled-coil sequence - N17, which flanks the polyQ tract and considerably enhances aggregation kinetics by promoting the formation of a critical tetrameric intermediate (4, 41). In contrast, the C-terminal proline-rich domain counteracts Httex1 aggregation by increasing the stability of oligomers (42). Therefore, it is essential to study how the N-terminal domain affects both fibrillation kinetics and resulting fibril morphology.

The Multi-eGO model assumes that the fibril state corresponds to the free energy minimum of the protein at high concentrations. While we have successfully developed a fibril structure for Q16, structural details and contact patterns of N17 domain in this fibrillar state remain largely unknown. Interestingly, previous studies indicated that N17 retains its α-helical structure within Httex1 fibrils (37). Thus, we conducted a dense phase simulation of the N17-Q16-P_5_ (H16) to mimic a crowded molecular environment and identify intermolecular contacts associated with the flanking N17/P_5_ domain. Specifically, we packed 170 chains of H16 into a cubic simulation box (concentration ∼100 mM, **Figure 4A**) and performed a 5 μs AAMD simulation. **Figure 4B** presents a comparison of the α-helical fraction of H16 dense phase with the dilute phase monomer, demonstrating an increased helicity within the N17 domain under the dense-phase conditions. Additionally, we observed the emergence of transient β-sheet and β-bridge structures in the dense-phase ensemble **(Figure 4C**). To inform the development of the multi-eGO model, we further analyzed the pairwise interchain contacts formed during the dense-phase simulation **(Figure 4D)**. The resulting two-dimensional contact map reveals numerous favorable interactions involving the flanking domains (N17-P_5_, N17-N17, and P_5_-P_5_),which considerably exceed those formed by the Q16 domain. Notably, interactions between N17 and Q_16_ domain were comparatively weaker.

**Figure 4:**
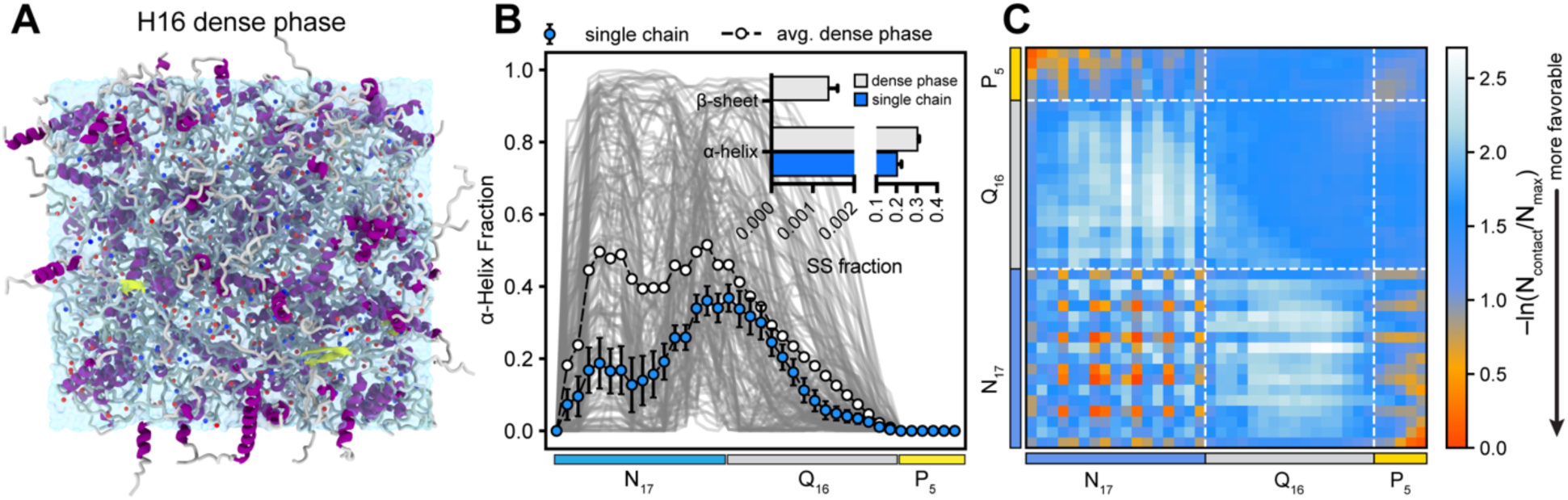
Structural properties of Q16-HttEx1 in the dense phase. (**A**) Snapshot illustrating 170 chains of Q16-HttEx1 in a cubic simulation box with a side length of 13 nm. Protein chains are represented as ribbons and Na^+^/Cl^−^ ions are depicted as blue and red spheres, respectively. (**B**) Average fractional helicity per residue, highlighting enhanced α-helical formation of the N17 domain within the dense phase compared to the dilute (single-chain) condition. Gray shading in the background represents helix profiles from individual protein chains. Inset shows the comparison of secondary structure (SS) fractions for H16 under dilute (single-chain) and dense-phase conditions, highlighting increased β-sheet formation in the dense phase. (**C**) Intermolecular vdW-based contact map, revealing substantial contributions of the N17 and P5 domains to multivalent interactions.

To study the H16 aggregation process, we developed the H16 Multi-eGO model by using the native contact information of the flanking domains from the dense-phase simulation. The contact information for the Q16 domain is taken from the semi-infinite simulations of fibril models as discussed in the previous sections **(Figure 2, 3)**. It is known that polyQ fibrils grow in both lateral and axial directions using hydrogen bonding and side chain interdigitation, respectively (43). Additionally, the cryoEM low resolution images show no interactions between flanking domains and the polyQ domain in the final aggregates (13). We, therefore, model the cross-interactions between the flanking domains and Q16 as excluded volume to avoid the hindrance in the growth of fibril and reduce the frustration on the energy landscape for the fibril growth. The contact information extracted from different ensembles for multi-eGO model of H16 is shown in **Figure S6**.

### Presence of flanking domains reduces fibril heterogeneities

Using the multi-eGO model of H16, we performed triplicate aggregation simulations of 500 chains at a concentration of 10 mM and a temperature of 300 K. Three independent aggregation simulations were performed for two H16 models parameterized using wither β-turn and β-arc configurations and each simulation was continued until almost all the chains were aggregated. Subsequently, we analyzed the evolution of the total β-sheet fraction **(Figure 5A)**.Notably, the two parameterizations led to markedly different aggregation behaviors. For the β-turn model, although H16 chains assembled into extensive aggregates, the Q16 domains within these assemblies remained largely disordered, yielding a low overall β-sheet fraction that plateaued at approximately 0.2 **(Figures 5A)**. In contrast, the using β-arc model promoted efficient β-sheet formation within the Q16 domains, resulting in a significantly higher β-sheet fraction of around 0.5 (**Figure 5A**). These structural differences reflected in the morphologies of the resulting aggregates (**Figure 5C**). Given these distinct results, and specifically because the β-arc parameterization was successful in producing β-sheet-rich H16 assemblies, all subsequent analyses and comparisons between Q16 and H16 models were carried out using the β-arc model.

**Figure 5:**
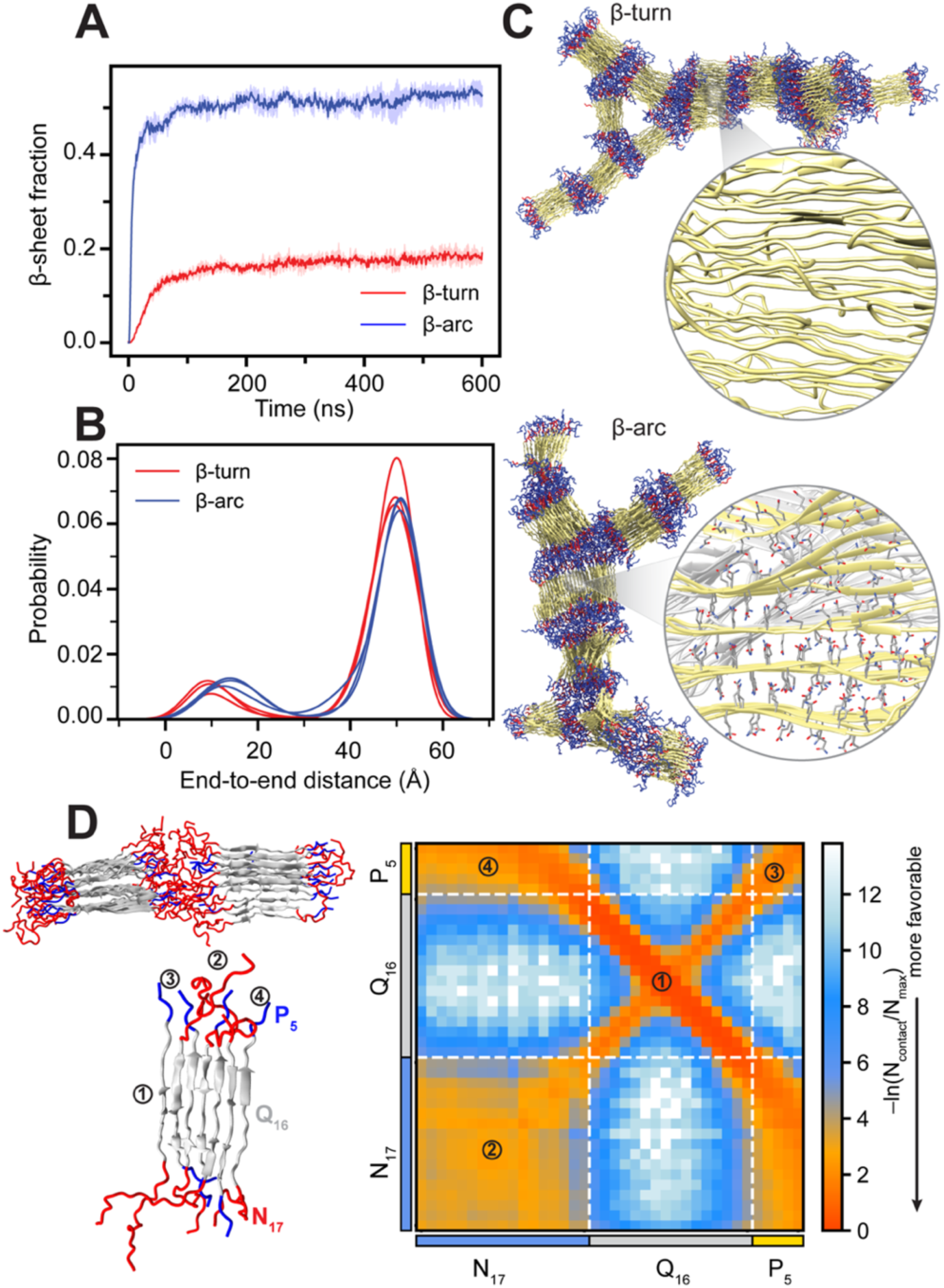
Structural properties of Q16-HttEx1 in multi-eGO aggregation simulations. (**A**) Time evolution of β-sheet fraction of Q16-HttEx1 β-turn and β-arc models simulated at 10 mM protein concentration and 300 K. End-to-end distance distributions for the two models, highlighting prevalence of extended conformations within fibril structures. (**C**) Representative snapshots of Q16-HttEx1 β-turn and β-arc fibril structures. Zoomed-in views illustrate the disordered assembly (β-turn) and extended layering and interdigitation of β-sheets (β-arc). (**D**) Intermolecular vdW-based contact map with snapshots highlighting predominant intermolecular interactions involving: (1) polyQ regions forming anti-parallel β-sheets, (2) N17-N17, (3) P5-P5, and (4) N17-P5 interactions.

To further investigate the conformational heterogeneity in the final fibril configuration associated with the presence of flanking domains, we compute the end-to-end distance distribution of the polyQ domain (aa:18-33) of H16. Our analysis indicates that individual chains strongly favor the β-strand (extended configuration) for the polyQ domain compared to β-arc or β-turn configurations with an overall reduction in conformational heterogeneity compared to Q16. In line with these observations, experiments indicate the short polyQ favors extended configuration while longer peptides begin to adopt turn configurations (40, 44) Additionally, the cryo-EM data suggests the flanking domain reduce the heterogeneities in the polyQ core and play an important role in stabilizing the polyQ core by preventing large out-of-register shifts Since the Q16 domain contact matrix in our H16 model is identical to the Q16 multi-eGO model, the reduced heterogeneity in the H16 multi-eGO fibril arises via flanking domain interactions.

We next analyzed the flanking domain interactions within the H16 fibril obtained from multi-eGO, presented as an interchain contact map in **Figure 5D**. This analysis reveals that the Q16 region adopts an antiparallel β-sheet configuration, while flanking domains exhibit considerably weaker interactions compared to the dense phase (**Figure 4C**). The N17 and P_5_ flanking domains arrange in an alternating pattern along both lateral and axial directions. Notably, each flanking domain is surrounded on all four sides by the other flanking domains, a feature that emerged spontaneously in the multi-eGO simulations and was not explicitly encoded in the contact matrix. At the supramolecular level, protofibrils interact through cross-contacts between flanking domains, consistent with the previous observations (7, 17). To further investigate the secondary structure content of different domains in the presence of explicit water and ions, we next perform all-atom simulations for protofibrils obtained from multi-eGO, which allows for an assessment of fibril stability under physiological conditions.

### Helical nature of N17 observed in AA simulations of multi-eGO protofibrils

Previous studies report that within Httex1 amyloid fibrils, N17 can adopt helical conformations, while the PRD remains largely disordered (5, 7, 45). Additionally, the polyQ fibril core is generally characterized as dehydrated (9). To assess the stability of H16 protofibrils obtained from multi-eGO simulations, we performed all-atom simulations in explicit solvent. Two different but stable protofibrils— comprising 19 and 22 chains, respectively—were selected from the H16 multi-eGo simulations, each representing aggregates formed in independent runs (**Movies S1, 2**). The representative structure of a solvated protofibril is shown in **Figure 6A** highlighting the characteristic side-chain interdigitation within the β-sheets and the dehydrated nature of the polyQ core. The time-dependent structural evolution of individual chains (**Figures 6B, S7C,D)** revealed stable β-sheet formation within the Q16 domain and the emergence of α-helical conformation of N17 domain. It is important to note that while N17 domains remain mostly disordered in the multi-eGO model, α-helical structures emerged only during the AAMD simulations (**Movies S1, 2**). This appearance of N17 helicity in the AAMD refinement likely underscores the importance of explicit solvent effects and detailed atomic interactions for stabilizing helical conformations within the protofibril context— particularly the specific attractive or guiding contacts between N17 and Q16 that were simplified to excluded volume in the multi-eGO fibril model.

**Figure 6:**
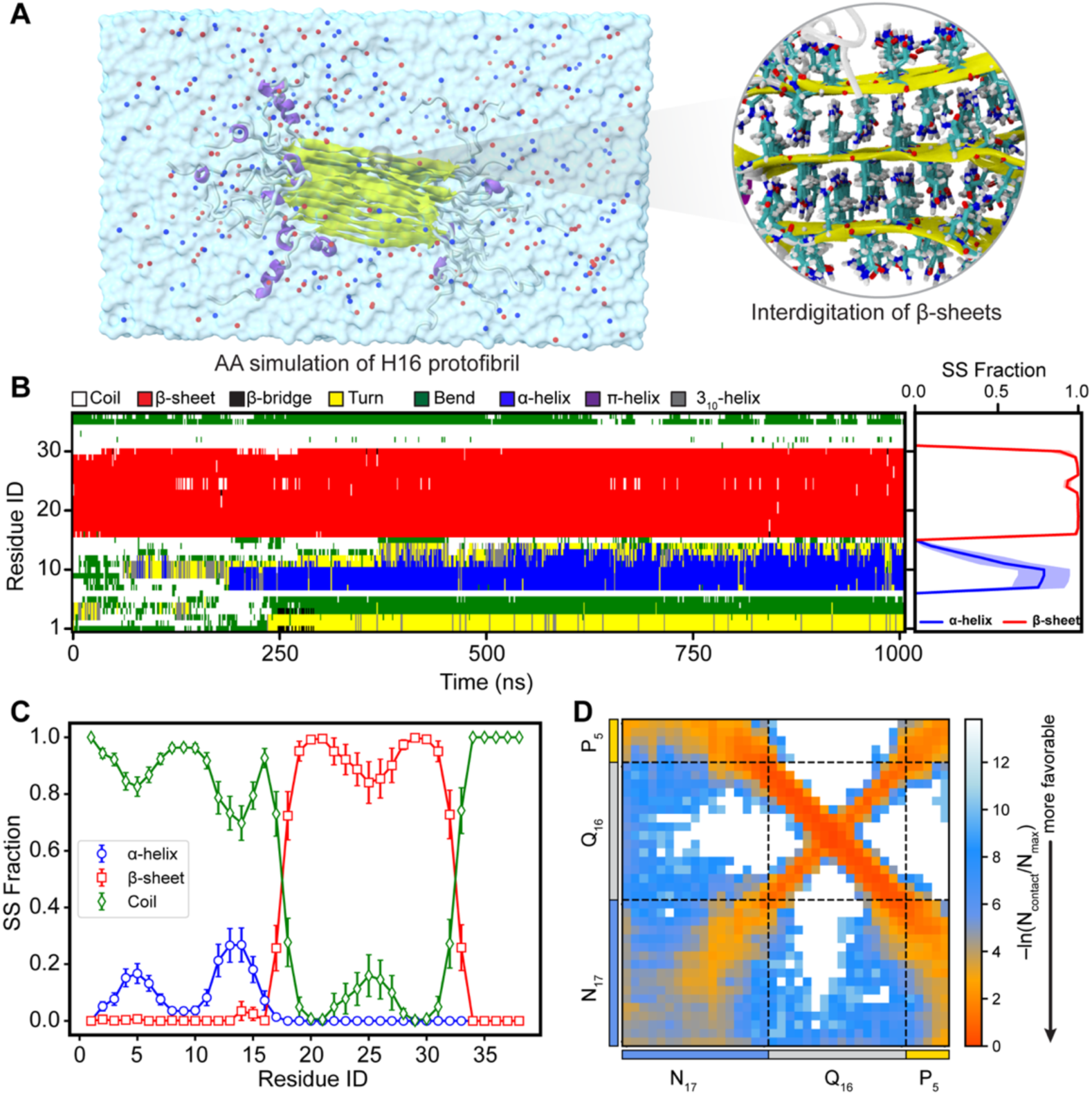
Conformational ensemble of a H16 protofibril. (**A**) Snapshot of the Q16-HttEx1 protofibril obtained from all-atom simulations. Protein chains are depicted as ribbons, Na^+^ and Cl^−^ ions as blue and red spheres, respectively, and water molecules as a transparent surface. A zoomed-in view highlights side-chain interdigitation within the fibril core. (**B**) Time-dependent secondary structure evolution of a representative protein chain, illustrating stable β-sheet structures in the polyQ fibril core and dynamic conformational changes of N17 region. Average fraction of secondary structure content across protofibril chains. Error bars represent standard deviations estimated by block averaging (four blocks). (**D**) Residue-based intermolecular vdW-based contact map within the protofibril, exhibiting interaction patterns consistent with multi-eGO aggregation simulations.

The average secondary structure content over all chains is shown in **Figure 6C**, indicating the stable fibril morphology. The helical content of the N17 domain which was enhanced in the dense phase (**Figure 4B**) reduces considerably in the protofibril state. We observe a ∼20% helical content for N17 domain, and the C-terminus (P_5_) remains disordered. We observed a bimodal distribution of the helicity in the N17 domain, wherein the helicity of N17 domain first reduces around at position 10 before further increasing to 20% at the 12^th^ position. A similar behavior was observed for other two protofibrils extracted from the multi-eGO fibril (**Figures S7A, B**). These observations agree with a recent secondary chemical shift analysis using solution NMR, wherein a similar bimodal distribution of the helicity was observed in the N17 domain of HttEx1 (31). A recent structural model of Q44-HttEx1 determined using an integrative approach combining ssNMR and MD simulation, also reported a similar flanking domain ensemble (15). Overall, the three protofibrils remain largely stable in AA simulations, implying that the multi-eGO protofibrils represent stable arrangements. The interchain contact map of a protofibril from AA simulations (**Figure 6D**) shows highly favorable residue pair interactions in the H16 fibril state due to its well-ordered and stable arrangement. Overall, all-atom simulations of protofibrils highlight the stability of multi-eGO fibrils, further validating the robustness of our multi-eGO modeling and simulation of Q16-Httex1 aggregation.

### Flanking domain (N17) enhances the aggregation kinetics by promoting the large order oligomers

To compare the kinetics and mechanism of Q16 and H16 aggregation, we performed multi-eGO simulations at submillimolar concentrations (1.0, 0.5, and 0.25 mM). Three independent aggregation simulations were performed, and the fraction of largest cluster size was analyzed as shown in **Figure S8**. The fraction of largest cluster is defined as the cluster with the maximum number of chains normalized with the number of chain present in the system. Comparison of the aggregation kinetics at 0.25 mM concentration indicates similar kinetics for both systems up to the first 25 ns of simulation **(Figure 7A)** after which the kinetics of H16 aggregation increases more rapidly than Q16 (**Movies S3, 4**). **Figure 7A** provides an illustration of different aggregation states, where Q16 forms highly heterogeneous aggregates (e.g., mixture of β-turn and extended stacking) compared to H16, which forms more structured fibrillar assemblies. H16 has a higher tendency to aggregate into structured fibrillar forms, possibly due to additional residues (N17 and P_5_), which enhance intermolecular interactions and increase structural order **(Figure 5B)**. Q16, on the other hand, aggregates slower and forms less structured aggregates with higher intra-fibril structural heterogeneities, indicating a weaker aggregation propensity. We observe a trend of increasingly faster kinetics for H16 at higher concentrations **(Figure S8)**, with the largest difference observed at 10 mM concentration. Overall, our models qualitatively capture the well-established enhancement in aggregation kinetics associated with the N17 flanking domain (4).

**Figure 7:**
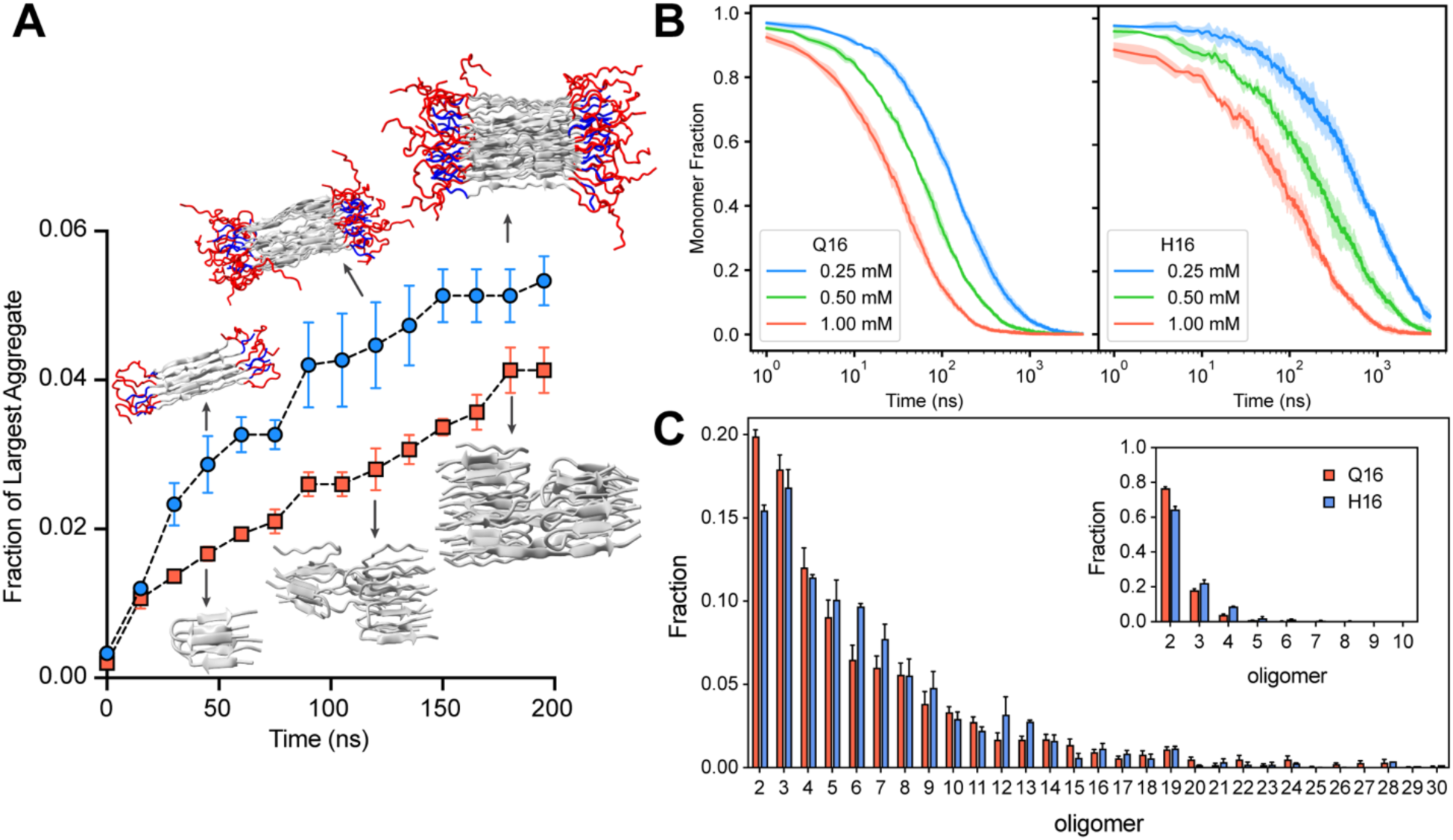
Aggregation kinetics of Q16 and H16. (**A**) Time evolution of the maximum cluster fraction for Q16 and H16 at 0.25 mM protein concentration and 300 K. Snapshots depict representative clusters formed during the simulations. (**B**) Fraction of monomers over time for Q16 and H16 at varying protein concentrations. (**C**) Oligomer order distributions before (inset) and after t1/2, the time at which the monomer fraction decreases by half. Error bars denote std. error of the mean, calculated over three independent trajectories.

To further probe into the origin of differences in the kinetics of H16 versus Q16 aggregation, we examined the monomer depletion rate **(Figure 7B)**. Q16 exhibits faster monomer depletion compared to H16, as seen by the faster decline in monomer fraction for all three concentrations. The faster depletion rate for Q16 suggests its monomers more readily engage in initial binding events, primarily forming smaller oligomers such as dimer. In contrast, the H16 monomer population decreases more gradually. Interestingly, while the growth of the largest aggregate over time suggests faster aggregation for H16 **(Figure 7A)**, the slower monomer depletion rate points to a more gradual initial kinetics. To better understand this discrepancy, we analyzed the oligomeric size distribution formed by both proteins before and after the monomer depletion half-life (*t_1/2_*), as shown in **Figure 7C**. Distributions were computed for up to 200 ns following *t_1/2_*. The oligomer size distribution indicates that small oligomers (dimers, trimers) are abundant early, with the percentage rapidly more decreasing for larger oligomer sizes. Before *t_1/2_*, the Q16 initially formed a larger fraction of dimers (∼75%) compared to H16 (∼65%).

However, the fraction of higher order oligomeric species (e.g., trimers, tetramers) are more abundant in the case of H16. Interestingly, H16 displays substantially higher percentages of mid-sized oligomers (oligomer size ∼5–13) compared to Q16, highlighting its greater propensity to form more oligomeric aggregates. The oligomeric size distribution computed after *t_1/2_* also shows a similar trend where the percentage of higher order oligomer (pentamers and beyond) is higher for the H16. We observe a similar trend for aggregation simulations at 0.5 and 1 mM protein concentrations (**Figure S9**).This behavior explains a higher monomer depletion rate for Q16 to form dimeric states that slowly converts into higher order oligomeric states than H16. We further computed the interchain contact map at different phases during the aggregation process, i.e., monomers, disordered clusters, ordered clusters, and protofibrils **(Figure S10)**. In the monomeric phase, the interchain contact map does not show β-sheet structure formation for either Q16 or H16. During the disordered cluster phase, both Q16 and H16 aggregation start to show the appearance of the antiparallel β-sheet configurations. Overall, H16 shows a greater tendency to form stable, mid-sized oligomeric aggregates, a critical intermediate in fibril formation (46, 47). In contrast, Q16 aggregates primarily into smaller oligomers, suggesting a limited ability to progress efficiently into larger stable structures.

In the early stages of aggregation observed in our simulations, individual chains transition from random coil states towards β-sheet structures. This process appears consistent with established principles where such conversions proceed through disordered intermediates (48–51). This structural conversion typically integrates with a “dock-and-lock” mechanism proposed for polyQ aggregation (52), where chains first associate (“dock”) with an existing oligomer or protofibril before conformationally rearranging (“locking”) into the fibril structure. Our detailed visual analysis of the Q16 simulations reveals two types of dock-and-lock mechanisms - (i) a backbone-mediated mechanism **(Figure S11A)** and (ii) a side-chain interlocking **(Figure S11B)** mechanism. In case of the backbone-mediated mechanism, a chain docks onto another chain through specific interchain backbone interactions followed by its conversion to a full β-sheet conformation. In the side-chain interlocking mechanism, an incoming chain docks primarily via side-chain interactions onto a template molecule (often one already in a β-turn). This initial association is followed by cooperative side-chain rearrangements between both chains, stabilizing the interaction through proper interdigitation and leading to the incorporation of the new chain, frequently resulting in a β-turn structure itself. A recent two bead per-residue coarse-grained model that applies a specific side-chain hydrophobic interaction potential to stabilize the polyQ fibrils and study polyQ aggregation also reported a similar two-step mechanism (53). For H16 aggregation, which includes N17 domain, previous studies suggest that the process is potentially driven by interactions involving either Q16 domain or N17 domain(4, 54) Our simulations provide insights into a plausible pathway for H16 assembly: initial association appears facilitated by interchain Q16 domain contacts that promote the alignment and formation of β-sheet structures (**Figure S11C**). Following this backbone ordering and β-sheet formation, these structured segments then dock laterally, stabilized by side-chain interactions, to contribute to the growing protofibril (**Figure S11D**).

## DISCUSSION

PolyQ and huntingtin amyloid fibrils are known to exhibit a high degree of structural polymorphism. Low-resolution cryoEM studies reveal substantial polymorphism in polyQ fibrils, which appears to be reduced upon addition of the N-terminal flanking sequence (13, 14). In contrast, ssNMR studies across different polyQ lengths, with and without flanking domains, suggest that this heterogeneity may not primarily originate at the secondary structure level of the fibril core (4, 5, 7). While a recent structural integrative approach has resolved a single, plausible structure of the Q44-HttEx1 (15), our multiscale simulations specifically identify polymorphism occurring at the tertiary and quaternary structural levels, as well as supramolecular organization of polyQ fibrils. To characterize polyQ domain polymorphism, we modeled two plausible tertiary structures—β-turn and β-arc—and arranged them in multiple directional configurations, resulting in different quaternary assemblies. These directional arrangements of Q16 monomers were consistent with experimental constraints from ssNMR and X-ray diffraction data, supporting the existence of polymorphism at the molecular level. Our results explicitly demonstrate that the cross β-strand architecture can be organized in multiple ways while still satisfying existing experimental constraints. Although our study primarily focused on directional arrangements of the termini, additional heterogeneity may arise from interactions between C- and N-termini of adjacent molecules, further expanding the structural diversity. Similar polymorphic arrangements of polyQ fibril have been reported in prior MD simulation studies (55). These results underscore the need for higher-resolution experimental techniques to resolve atomic-level polyQ fibril structure.

Our multi-eGO aggregation simulations of Q16 offer important insights into amyloid fibril structure and its inherent heterogeneity. The resulting fibrils exhibit a variable-width, branched morphology, closely resembling structures observed in atomic-forced microscopy and negative stain transmission electron microscopy studies (4, 11). Detailed structural analysis of the fibril architecture reveals substantial heterogeneity at the tertiary structure level, with individual chains adopting diverse conformations including β-turn, β-arc, and β-strand arrangements. This structural diversity suggests a potentially general characteristic for amyloid assemblies formed by other homopolymeric sequences, such as polyalanine and polyproline. Notably, our simulations also capture the formation of small oligomeric species adopting β-strand conformation, reminiscent of pathogenic β-hairpin structures reported for polyQ (56) and HttEx1 (57) in their monomeric and early oligomeric species. Given that small protofibrils have been implicated more cytotoxic than mature amyloid fibrils (17), these findings underscore the importance of characterizing early-stage fibrillation events in greater details. Mechanistically, our simulations reveal two distinct dock-and-lock mechanisms for polyQ aggregation and highlight the critical role of the partially helical N-terminal flanking domain in accelerating H16 fibrillation. Specifically, the presence of N17 facilitates the formation of large, ordered oligomers and enhances aggregation kinetics. The interplay between helical content and conformational flexibility is increasingly recognized as a key factor modulating both protein condensates and conformational transitions (49, 50, 58). Accordingly, the enhanced α-helical of the N17 domain observed in the dense phase may present a key driver of pathological aggregation. Overall, our study demonstrate the power of multiscale simulation to dissect early-stage fibrillation and provide detailed molecular perspectives on amyloid fibril structure, polymorphism, and the influence of flanking domains.

Our comprehensive investigation of H16, combining multi-eGO aggregation simulations with AAMD refinement of protofibrils, reveals several key findings that both validate and expand upon previous studies. The multi-eGO simulations demonstrate that flanking domains in H16 enhance aggregation kinetics compared to isolated Q16, consistent with experimental observations (37). These flanking regions also promote the formation of amyloid fibrils with reduced structural heterogeneity, favoring predominant β-strand configurations. Subsequent AAMD simulations confirm the structural stability of these H16 protofibrils while providing detailed insights into their secondary structural elements. Our analysis reveals that the N17 domain adopts a partially α-helical conformation (∼20% helical content), addressing the range of structural interpretations presented in previous literature. Simultaneously, the C-terminal P5 domain maintains its disordered and flexible nature, aligning with earlier reports (37, 59, 60). These findings highlight how specific characteristics of the flanking domains influence both aggregation dynamics and the resulting fibril architecture. Our study demonstrates the power of multiscale simulations to explore the fibrillation process across large spatiotemporal scales, providing molecular insights into early aggregation events—critical stages implicated in many neurodegenerative diseases. This work also makes a significant contribution toward characterizing the structural polymorphism of polyQ fibrils and proposing plausible molecular architectures for Huntingtin amyloid fibrils. Specifically, by employing a minimal polyQ repeat model, we demonstrate that polyQ fibrils can adopt diverse tertiary and quaternary structures, suggesting that such polymorphism may be an intrinsic feature of aggregation-prone homopolymeric domains. Given that fibril polymorphism can profoundly influence drug binding and efficacy, these molecular insights offer a valuable framework for designing more effective therapeutic strategies targeting early-stage aggregation in polyQ-associated neurodegenerative disorders.

## MATERIALS AND METHODS

### All-atom simulation of monomer and fibril state of Q16 and H16

The initial monomer coil configurations of Q16 and Q16-HttEx1 were generated using MODELLER (61). For the fibril structure of the Q16, we use two proposed tertiary antiparallel β-sheet structures of Q16 fibril, i.e., β-turn and β-arc (**Figures 2A,E**) and stack them in a total of eight different end-terminus directional arrangements as shown in **Figure S2**, while maintaining the experimentally observed sheet-to-sheet and strand-to-strand distances (5, 62, 63). All eight arrangements were put in a semi-infinite setup where the fibrils were allowed to interact with their periodic image in X and Y directions as shown in **Figures 2B,F**. Q16-HttEx1 protofibrils were taken from the output of Multi-eGO simulations. The description of multi-eGO simulations is provided in the next section. To prepare the all-atom (AA) Q16-HttEx1 dense phase simulation, we use a method similar to the one described previously (64). Briefly, to obtain the initial structure for multichain simulation, 170 chains of the Q16-HttEx1 were first equilibrated at 300 K using coarse-grained (CG) simulation with HPS-SS model (65). The AA dense phase configuration was reconstituted from the Cα positions using the CG configuration using Modeler. Potential steric clashes were resolved with by running short implicit solvent simulations using AMBER03 force field (66)and OBC implicit solvent model (67) in OpenMM 8.1.1 (68).

All the initial configurations above were solvated in a cubic box with 150 mM NaCl salt concentration to mimic the physiological condition. Additional counter ions were added to maintain the charge neutrality. The force field parameters from protein were taken from AMBER03ws force field (https://bitbucket.org/jeetain/all-atom_ff_refinements/src/master/) (69), TIP4P/2005 model was used for solvent (70), and modified LJ parameters were used for Na^+^ and Cl^-^ ions.(71). The solvated structures were first energy minimized using steepest descent minimization algorithm in GROMACS-2020.4. The energy minimized structures were then thermally equilibrated at 300 K in a canonical (NVT) ensemble using Nosé–Hoover thermostat (72) with a coupling constant of 1.0 ps. Further, NPT equilibrations were carried out using Berendsen barostat with semi-isotropic pressure coupling for semi-infinite Q16 fibril simulations and isotropic pressure coupling for the rest of the systems, and a coupling constant of 5.0 ps was used to maintain the pressure at 1 bar (73). Final production runs in NPT ensemble were performed using OpenMM 8.1.1. We use the hydrogen mass repartitioning scheme by setting the hydrogen mass to 1.5 amu to use a larger 4 fs time step (74) Long-range electrostatic interactions were calculated using the particle mesh Ewald (PME) method (75). The short-range van der Walls interaction was set to 0.9 nm. The SHAKE algorithm was used to apply constraints to all hydrogen bonds (76).

### Multi-eGO model development and simulation details of Q16 and H16

The multi-eGO models of Q16 and Q16-HttEx1 were prepared using the previously described methodology (27, 77). The multi-eGO model is an implicit solvent model with a united atom representation of the system, where the parameters for bond, angle, and dihedral potential are transferable and taken from the GROMOS 54A7 force field. The non-bonded native contacts are learned from the different sources as described in the main text: all-atom monomeric ensemble, all-atom dense phase simulations and amyloid all-atom ensemble. The non-native contacts are modeled as excluded volume using basic Multi-eGO repulsive interactions. The free parameter of the Multi-eGO model, the interaction strength of native contacts (ε) for Q16 and Q16-HttEx1 were optimized by reproducing the monomer ensemble of the respective all-atom simulations (**Figure S1**).

Following the Multi-eGO simulation protocol (25), the Q16 and Q16-Httex1 systems modeled using the Multi-eGO model was minimized using Gromacs-2020.4 using the steepest descent algorithm until maximum force converges to value < 1000 kJ mol^−1^nm^−1^, followed by conjugate-gradient minimization until maximum force converges to value < 10 kJ mol^−1^nm^−1^. Next, position restrained relaxation was performed at 300 K in NVT ensemble. The short-range vdW cutoff was used as 1.45 nm, and a 5 fs time step was used in the simulations. The GPU-accelerated production simulations of Multi-eGO models were performed in the NVT ensemble at 300 K, with Langevin middle integrator, the friction coefficient of 1 ps^−1^, using the OpenMM 8.1.1.

### Trajectory analysis and visualization

Secondary structure content was computed using DSSP algorithm (78, 79) integrated in GROMACS. The contact maps, dihedral angles, and interatomic distances were analyzed using in-house scripts built on the MDAnalysis (80) package (version 2.5.0). Clustering analysis was performed based on Cα-Cα distance between two molecules, using a 10 Å cutoff to identify nearest neighbors. Pairwise residue contacts were defined as van der Waals interactions if at least one heavy atom from a residue was within 6 Å of a heavy atom from another residue. This distance threshold, validated in several previous studies (18, 81), reliably captures a range of interaction types including van der Waals forces, hydrogen bonds, and salt bridges. All the visualizations were rendered using VMD (82) and Chimera/ChimeraX (83, 84) visualization software.

## AUTHOR INFORMATION

### Corresponding Authors

*

*

*

*

*

### Author Contributions

AK: Multi-eGO Model parameterization, MD simulations, post-processing, data curation, interpretation, writing – original draft, TMP: All-atom dense phase simulation, data presentation, visualization, analysis, interpretation, writing – review and editing, AR: setting up the workflow of multi-eGO simulations, review and editing, PM: project administration, writing – original draft, review and editing, JM: funding acquisition, conceptualization, project administration, supervision, writing – review and editing.

## ACKNOWLEDGMENT

The presented work is supported by the National Institutes of Health under the grant R35GM153388. We thank Randal Halfmann from Stowers Institute of Medical Research for providing the initial single-unit structure of β-turn and β-arc structures. The authors acknowledge the Texas A&M High Performance Research Computing (HPRC) for providing computational resources that have contributed to the results reported in the article. We thank Carlo Camilloni for making the scripts and parameters of the m ulti-eGO force field available on GitHub.

