## Supplemental Information for "Multiscale simulations elucidate the mechanism of polyglutamine aggregation and the role of flanking domains in fibril polymorphism"

### Supporting Information

#### 1. Supporting Table.

**Table S1**: The comprehensive details of the simulations performed in our study.

| System information | Initial (final state) | Number of chains | Simulation time (ns) |
|---|---|---|---|
| Q16 monomer | IDP (partial helix) | 1 | 1000 |
| All-atom Q16 (semi-infinite) | BT-A1 fibril (fibril) | 16 | 1000 |
| | BT-A2 fibril (fibril) | 16 | 1000 |
| | BT-A3 fibril (fibril) | 16 | 1000 |
| | BT-A4 fibril (fibril) | 16 | 1000 |
| | BA-A1 fibril (fibril) | 16 | 1000 |
| | BA-A2 fibril (fibril) | 16 | 1000 |
| | BA-A3 fibril (fibril) | 16 | 1000 |
| | BA-A4 fibril (fibril) | 16 | 1000 |
| Multi-eGO (10 mM Q16) | BT-A1 dilute (fibril) | 1000 | 1000 × 3 |
| | BA-A1 dilute (fibril) | 1000 | 1000 × 3 |
| Multi-eGO (1 mM Q16) | BT-A1 dilute (fibril) | 1000 | 1000 × 3 |
| | BA-A1 dilute (fibril) | 1000 | 1000 × 3 |
| Multi-eGO (0.5 mM Q16) | BT-A1 dilute (fibril) | 1000 | 1000 × 3 |
| | BA-A1 dilute (fibril) | 1000 | 1000 × 3 |
| Multi-eGO (0.25 mM Q16) | BT-A1 dilute (fibril) | 1000 | 1000 × 3 |
| | BA-A1 dilute (fibril) | 1000 | 1000 × 3 |
| Q16-HttEx1 monomer | IDP (partial helix) | 1 | 1000 |
| Q16-HttEx1 all-atom dense phase | Dense phase (partial helix) | 170 | 5000 |
| Multi-eGO (10 mM Q16-HttEx1) | BT-A1 dilute (fibril) | 500 | 1000 × 3 |
| | BA-A1 dilute (fibril) | 500 | 1000 × 3 |
| | BT-A1 dilute (fibril) | 500 | 1000 × 3 |

| | | | |
|---|---|---|---|
| Multi-eGO (8 mM Q16-HttEx1) | BA-A1 dilute (fibril) | 500 | 1000 × 3 |
| Multi-eGO (6 mM Q16-HttEx1) | BT-A1 dilute (fibril) | 500 | 1000 × 3 |
| | BA-A1 dilute (fibril) | 500 | 1000 × 3 |
| Multi-eGO (4 mM Q16-HttEx1) | BT-A1 dilute (fibril) | 500 | 1000 × 3 |
| | BA-A1 dilute (fibril) | 500 | 1000 × 3 |
| Multi-eGO (2 mM Q16-HttEx1) | BT-A1 dilute (fibril) | 500 | 1000 × 3 |
| | BA-A1 dilute (fibril) | 500 | 1000 × 3 |
| Multi-eGO (1 mM Q16-HttEx1) | BT-A1 dilute (fibril) | 500 | 1000 × 3 |
| | BA-A1 dilute (fibril) | 500 | 1000 × 3 |
| Multi-eGO (0.5 mM Q16-HttEx1) | BT-A1 dilute (fibril) | 500 | 1000 × 3 |
| | BA-A1 dilute (fibril) | 500 | 1000 × 3 |
| Multi-eGO (0.25 mM Q16-HttEx1) | BT-A1 dilute (fibril) | 500 | 1000 × 3 |
| | BA-A1 dilute (fibril) | 500 | 1000 × 3 |
| All-atom simulation of Q16-HttEx1 protofibril | Fibril (fibril) | 19 | 1000 |
| | Fibril (fibril) | 22 | 1000 |

#### 2. Supporting Movies

**Movie S1**. All-atom molecular dynamics (AAMD) simulation of an H16 protofibril composed of 19 chains. The structure is color-coded by secondary structure: α-helix (purple), β-sheet (yellow), and coil/turn/bend (silver).

**Movie S2**. AAMD simulation of an H16 protofibril composed of 22 chains. The structure is color-coded by secondary structure: α-helix (purple), β-sheet (yellow), and coil/turn/bend (silver).

**Movie S3**. Multi-eGO simulation of a Q16 system containing 1000 chains. Chains are color-coded individually to distinguish different molecules.

**Movie S4**. Multi-eGO simulation of an H16 system containing 500 chains. Domains are color-coded as follows: N17 = red, Q16 = silver, and P5 = blue.

#### 3. Supporting Figures

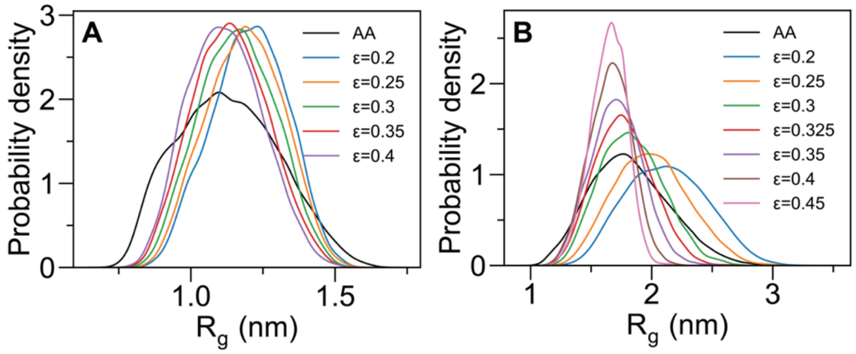


**Figure S1**: **Parameterization of Multi-eGO models against the AA monomeric ensemble**. Radius of gyration ($R_g$) distribution for varying values of the free parameter (ε) in Multi-eGO simulation of (**A**) the Q16 peptide and (**B**) the Q16-HttEx1 protein, compared to the corresponding AA simulations.

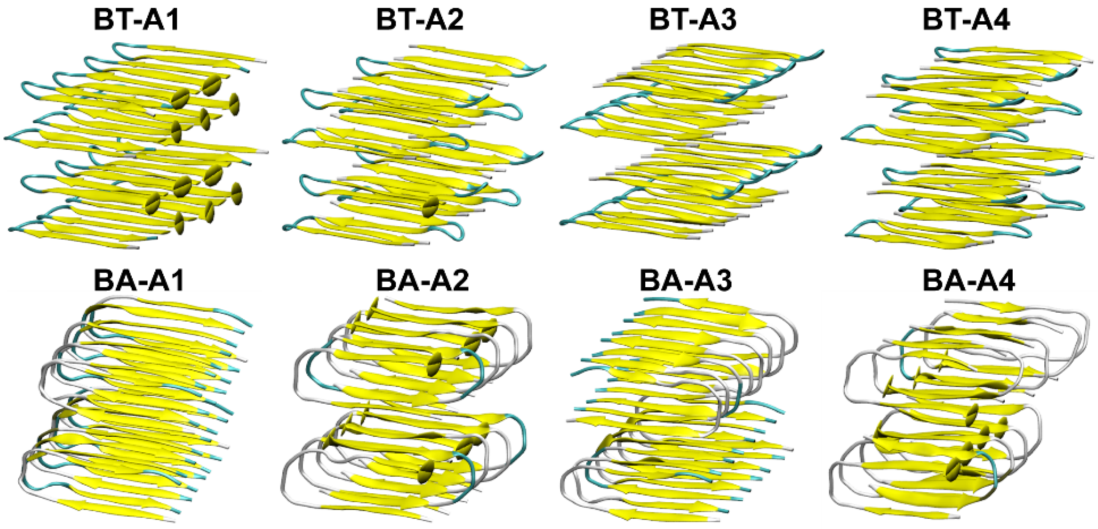


**Figure S2**: **Fibril arrangements considered in the study**. BT-A1: β-turn with the end-termini aligned in the same direction. BT-A2: β-turn with the end-termini arranged alternately along the lateral direction. BT-A3: β-turn with the end-termini arranged alternately along the axial direction. BT-A4: β-turn with the end-termini arranged alternately in both lateral and axial directions. Similarly, BA-A1, BA-A2, BA-A3, and BA-A4 represent the corresponding directional arrangements for the β-arc polyQ model.

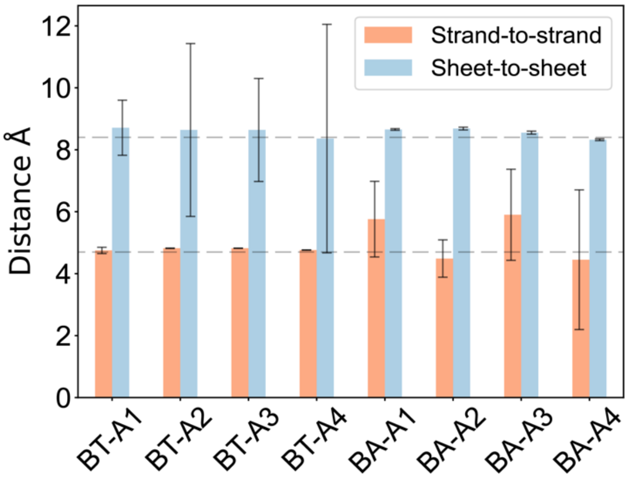


**Figure S3**: **Distance analysis of fibril arrangements**. Sheet-to-sheet and strand-to-strand distances for all directional arrangements considered in this study (Figure S2). The dashed lines represent distances measured from X-ray diffraction experiments: 4.75 Å for strand-to-strand spacing and 8.3 Å for sheet-to-sheet spacing.(6)

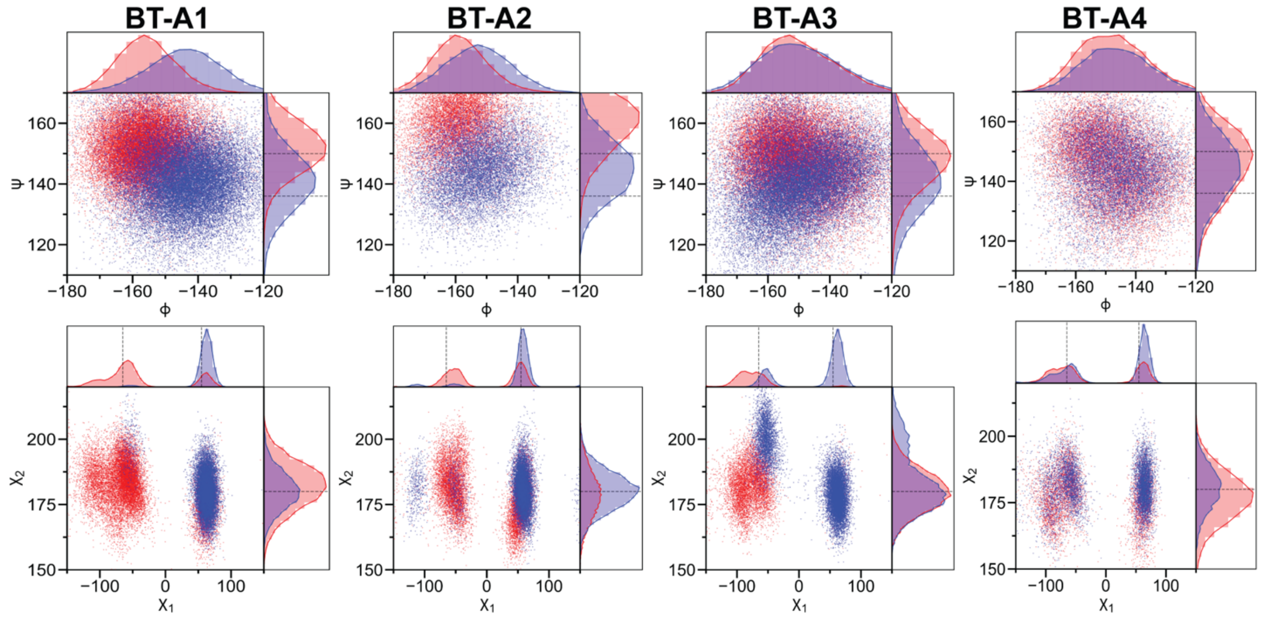


**Figure S4**: **Dihedral angle distributions of β-turn fibril arrangements**. Backbone (ϕ and ψ) and Side chain (χ1 and χ2) dihedral angle distributions for β-turn configurations shown in Figure S2. Black dashed lines indicate values observed in ssNMR experiments.(6)

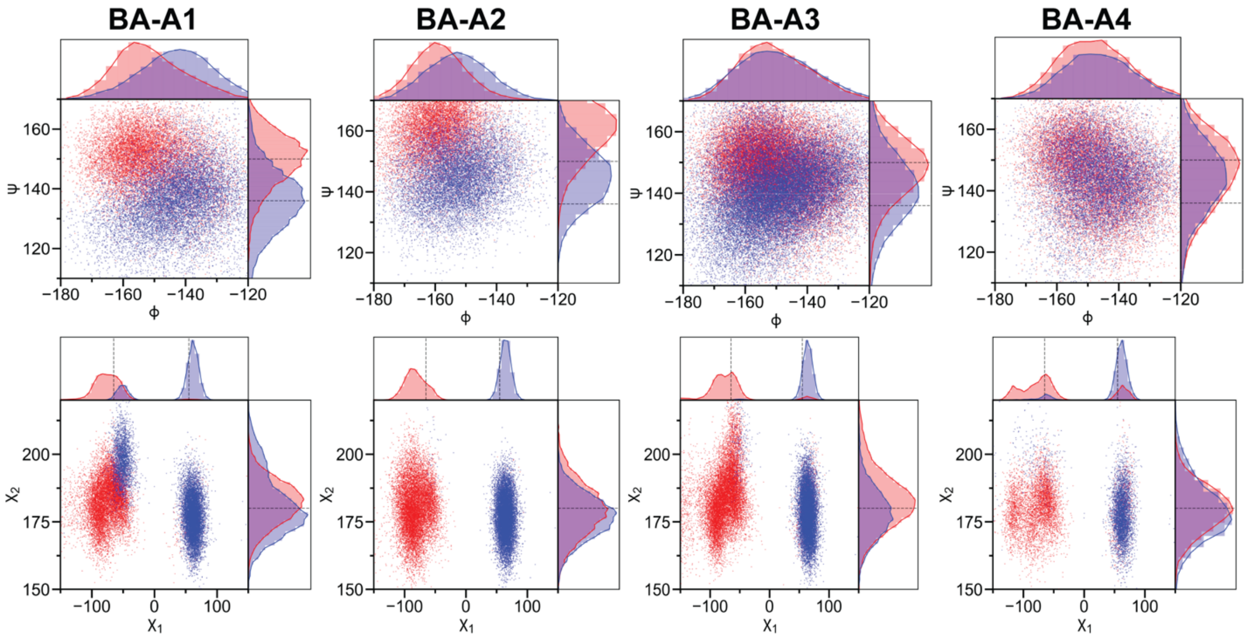


**Figure S5**: **Dihedral angle distributions of β-arc fibril arrangements**. Backbone (ϕ and ψ) and Side chain (χ1 and χ2) dihedral angle distributions for β-arc configurations shown in Figure S2. Black dashed lines indicate values observed in ssNMR experiments.(6)

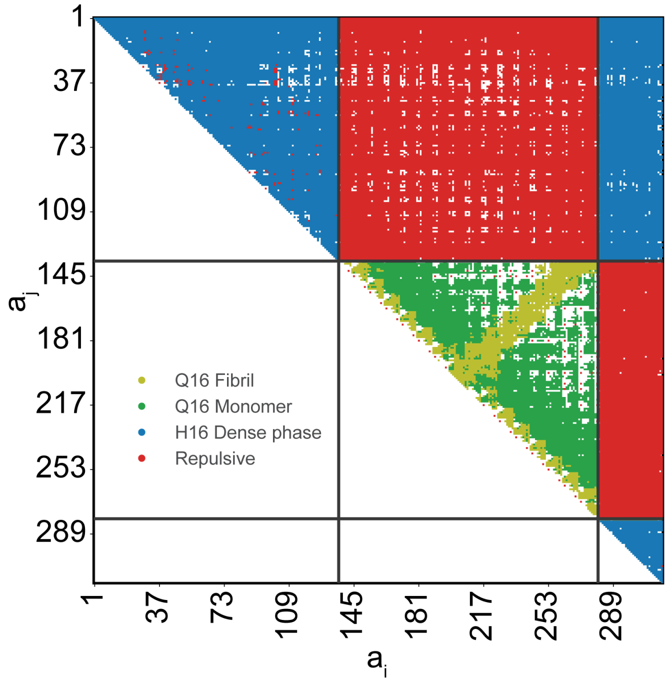


**Figure S6**: **Contact matrix for Q16-HttEx1 Multi-eGO Model.** All heavy atoms Multi-eGO Q16-HttEx1 model contact matrix incorporates contacts for the N17 and P5 flanking domains derived from the dense phase, while the contacts for Q16 are based on the Q16 Multi-eGO model. Interactions between the flanking domains and Q16 are designed to be purely repulsive to reduce frustration in the landscape.

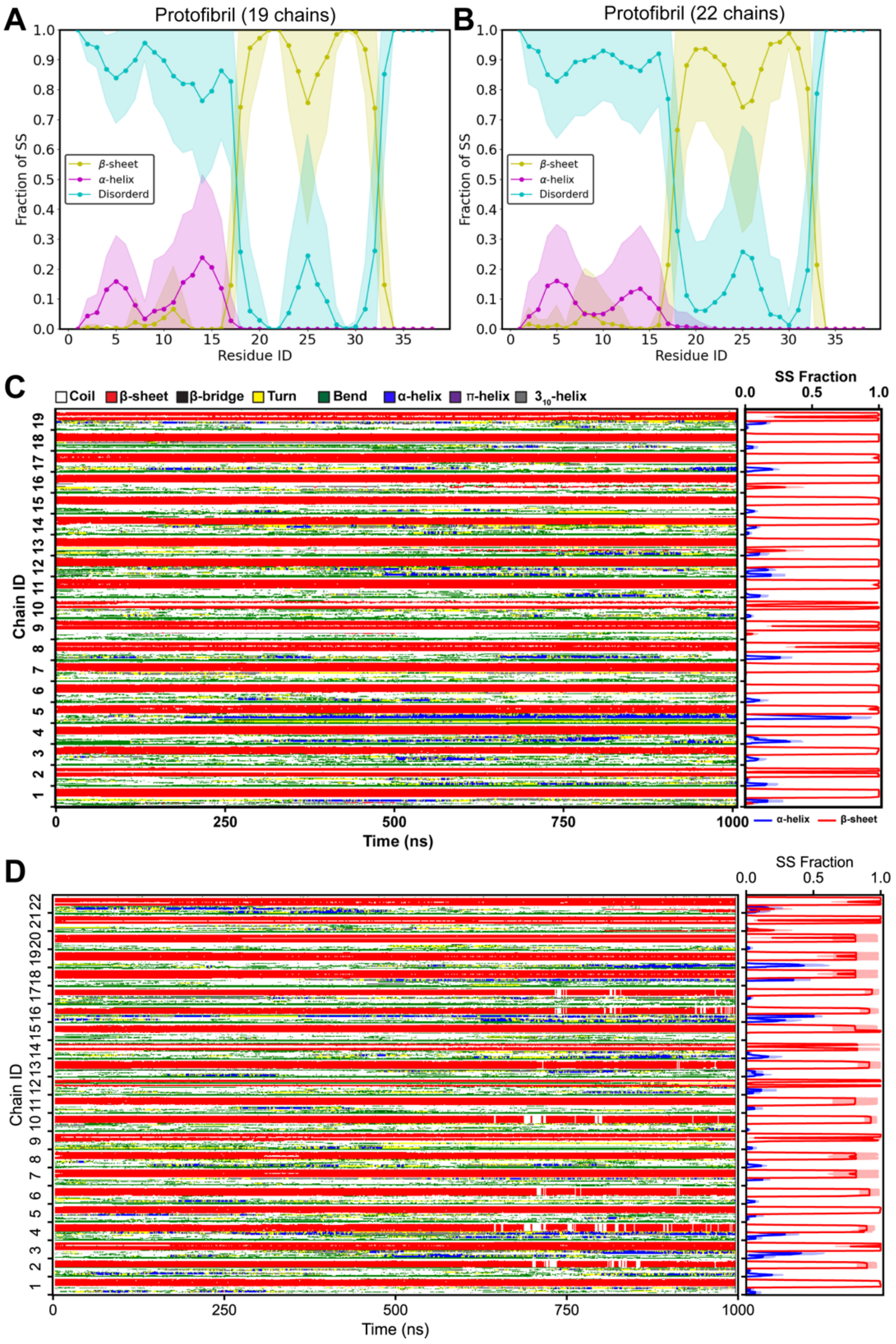


**Figure S7**: **Structural Characterization of Q16-HttEx1 Protofibrils from All-Atom Simulations.** (**A, B**) The average fraction of secondary structure content for tow selected protofibrils of different sizes (19 and 22 chains, respectively), with the shaded region representing the standard deviation. (**C, D**) Time-dependent secondary structure variation of all chains in the two protofibrils containing (**C**) 19 chains and (**D**) 22 chains, demonstrating stable β-sheet structures in the polyQ core and dynamic conformational changes in the N17 and P5 regions.

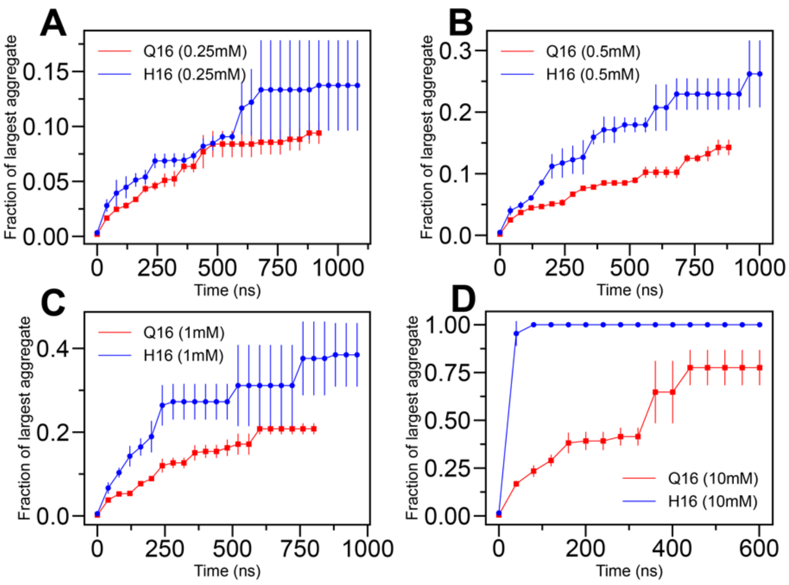


**Figure S8**: **Aggregation kinetics of Q16 and Q16-HttEx1.** Time evolution of the maximum cluster fraction of Q16 and Q16-HttEx1 at (**A**) 0.25 mM, (**B**) 0.5 mM, (**C**) 1 mM, and (**D**) 10 mM concentrations and 300 K temperature.

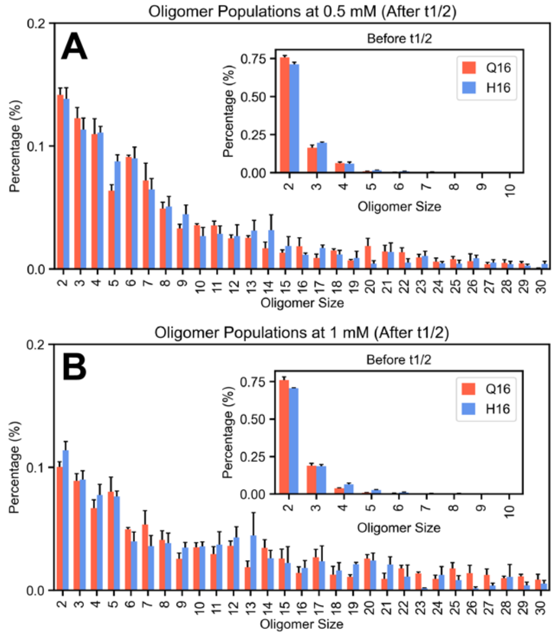


**Figure S9**: **Oligomer size distribution.** The oligomeric size distribution change before (inset) and after the half-life of monomer, the time at which the monomer fraction reduces by half, for (**A**) 0.5 mM and (B) 1 mM concentrations.

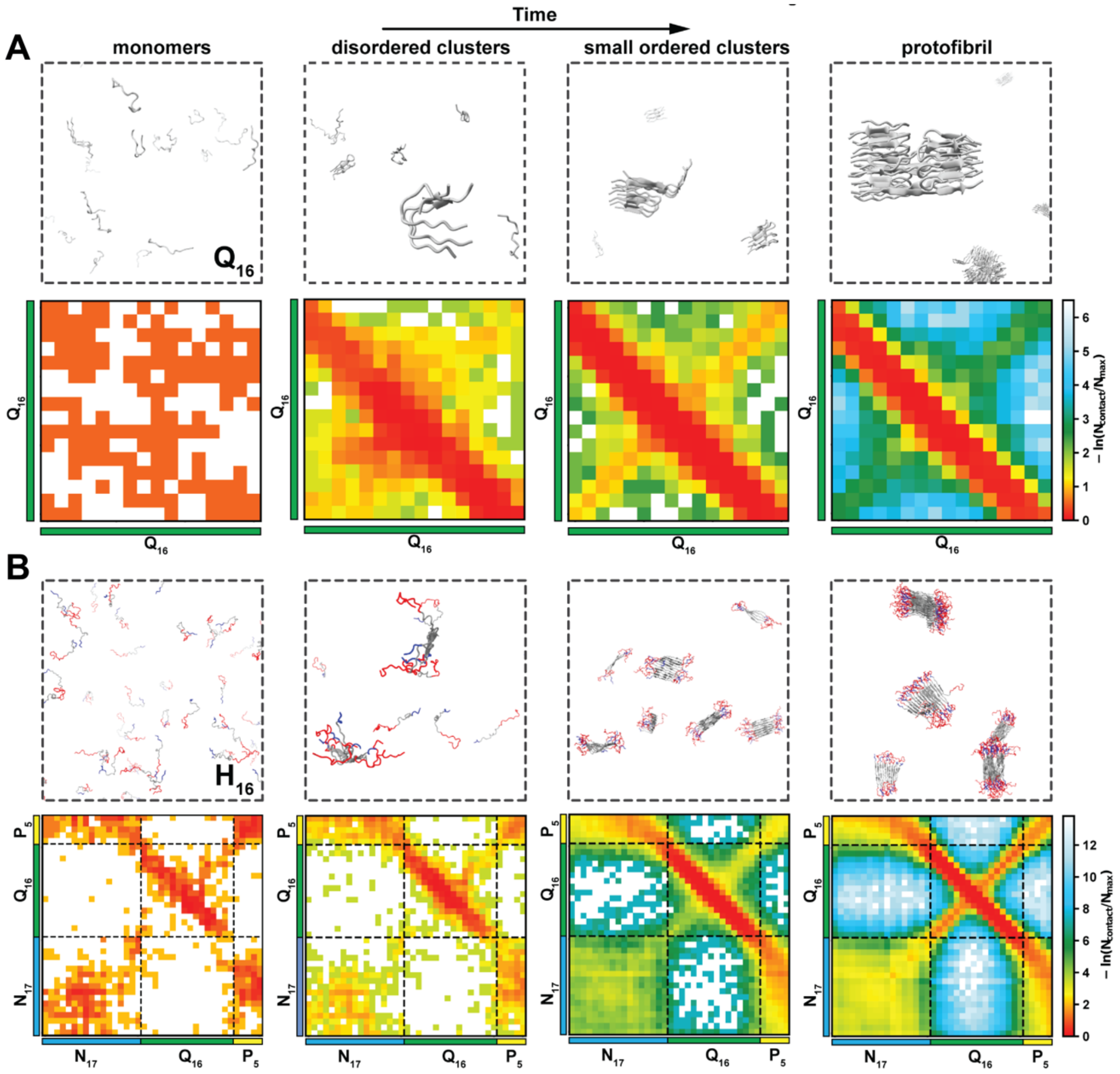


**Figure S10**: **Aggregation mechanism of Q16 and Q16-HttEx1.** Schematic representation the aggregation processes and evolution of intermolecular contacts for (**A**) Q16 and (**B**) Q16-HttEx1. Initially, monomers self-assemble into disordered clusters, which subsequently undergo structural rearrangements to form small, ordered clusters. These clusters serve as templates for further elongation, ultimately leading to protofibril formation.

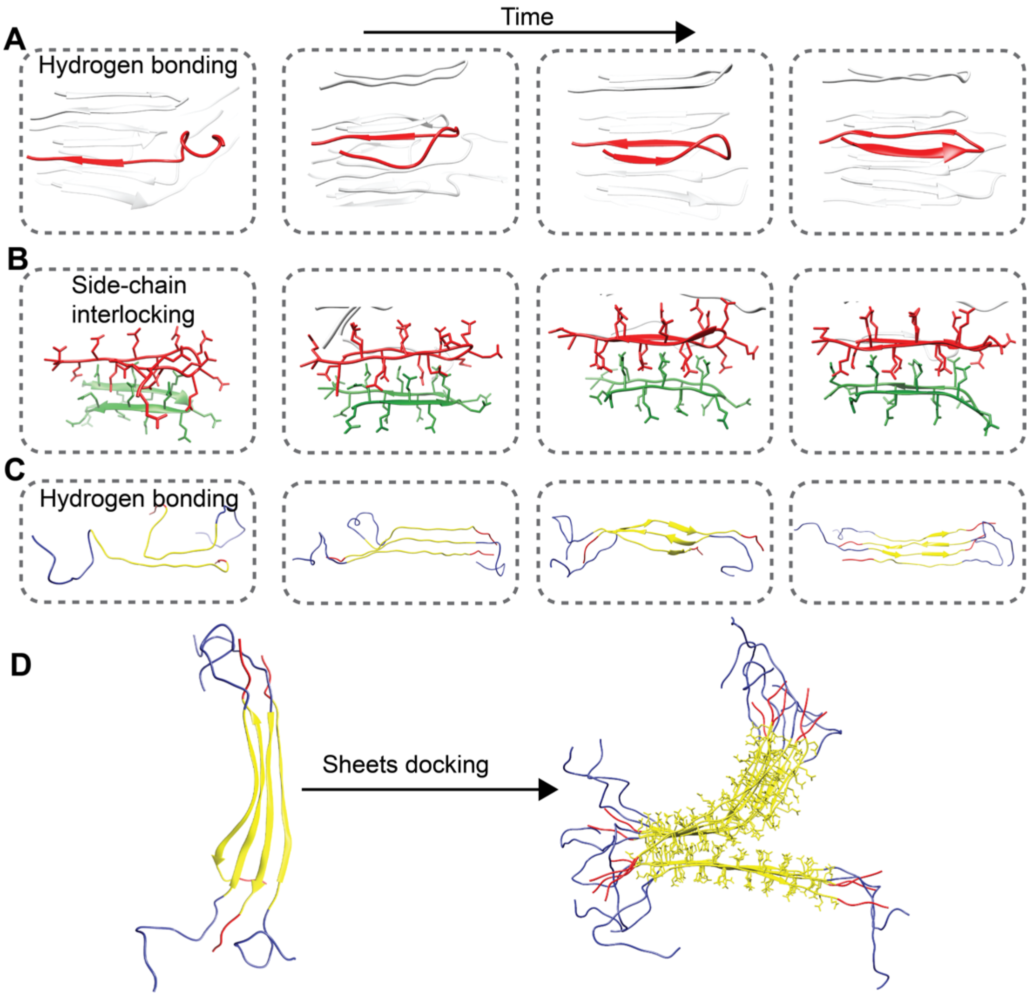


**Figure S11**: **Early stage aggregation mechanism of Q16 and Q16-HttEx1.** Two different types of dock-and-lock mechanisms observed in Q16 aggregation simulations – (**A**) Hydrogen bonding driven – the Q16 chain (red) docks through interchain hydrogen bonding, followed by full conversion to a β-sheet; and (**B**) Side-chain interlocking driven – chain-1 (green) initially exists in a β-turn conformation and serves as a template for chain-2 (red) to dock via its side chains. Once chain-2 fully docks, the side chains of both chains rearrange cooperatively, causing chain-1 to lose its β-turn structure. Ultimately, this cooperative motion allows both molecules to achieve proper side-chain interdigitation in a β-turn configuration. (**C**) Q16-HttEx1 aggregation is initiated by Q16 domain (yellow) interchain interaction, leading to the formation of β-sheets. (**D**) Subsequently, these β-sheets dock onto other β-sheets through side-chain interlocking to produce a protofibril.
